# The use of social information in vulture flight decisions

**DOI:** 10.1101/2023.07.26.550671

**Authors:** Sassi Yohan, Nouzières Basile, Scacco Martina, Tremblay Yann, Duriez Olivier, Robira Benjamin

## Abstract

Animals rely on a balance of personal and social information to decide when and where to move next in order to access a desired resource, such as food. The benefits from cueing on conspecifics to reduce uncertainty about resources availability can be rapidly overcome by the risks of within-group competition, often exacerbated toward low-ranked individuals. Being obligate soarers, relying on thermal updrafts to search for carcasses around which competition can be fierce, vultures represent ideal models to investigate the balance between personal and social information during foraging movements. Linking dominance hierarchy, social affinities and meteorological conditions to movement decisions of eight captive vultures, *Gyps spp*., released for free flights in natural-like soaring conditions, we found that they relied on social information (i.e. other vultures using/having used the thermals) to find the next thermal updraft, especially in unfavourable flight conditions. Low-ranked individuals were more likely to disregard social cues when deciding where to go next, possibly to minimise the competitive risk of social aggregation. These results exemplify the architecture of decision-making during flight in social birds. It suggests that the environmental context, the context of risk and the social system as a whole calibrate the balance between personal and social information use.

## 1. Introduction

Animals must constantly decide where and when to move next in order to find resources such as food, water, shelter, or a mate, necessary for life. To make these decisions, they can rely on two sources of information: personal information and social information. Personal information includes knowledge of the spatiotemporal patterns of resource distribution that individuals may perceive or have memorised from previous encounters [1]. For example, food-storing birds are able to return to locations where they stored or saw food in the past, based on prior expectation of the resource availability [2]. Social information, on the other hand, is obtained by observing the behaviour of others [3–5]. Feeding, fleeing, or mating individuals provide discrete information about the availability and locations of food, predators, or potential mates.

For resources that are heterogeneously distributed in the environment, ephemeral and unpredictable, using only personal information for movement decisions may be prone to inaccuracies [6]. In such conditions, social animals may benefit from companions’ knowledge and may follow the dominant or oldest individual(s) considered as knowledgeable (e.g. homing pigeons, *Columba livia*, or elephants, *Loxodonta africana*, [7,8]), follow the largest group through shared decision-making [9], or stay with preferred affiliates [10–12]. Because using social information can considerably reduce uncertainty in finding resources, individuals should favour this source of information to achieve cost-efficient movement [13–15]. However, relying heavily on social information can also lead individuals to aggregate on resources, potentially inducing competition by exploitation or interference if the resource is monopolizable and depletable [16]. Since both social and personal information are often available to social animals [1], they need to balance their relative importance, depending on the availability and predictability of the resource. When deciding on the next movement step, social animals must trade-off the decreased uncertainty of locating a resource through social information, with the potential increase in competition risk. Such a balance may be dictated by the immediate needs of the individual and its risk sensitivity [17] but also by the group social organisation. For example, low-ranked individuals are known to suffer more from within-group competition compared to high-ranked individuals [18] and should therefore be more reluctant to engage into social information use, which could eventually trigger proximity to despotic individuals [19].

Vultures rely on two unpredictable resources: carcasses to feed and thermal updrafts to move. During foraging flights, these large soaring birds gain altitude by circling into thermal updrafts (i.e. masses of hot air rising from heated surfaces) and glide across the landscape to the next updraft while scanning the ground for carcasses [20]. Although some topographic features are clearly favourable to updrafts presence [21], at the individual level, challenging local meteorological conditions (e.g. high wind speed, low temperature, high cloudiness) can make thermal locations and availability hard to predict [22]. If they fail to detect an updraft, vultures may be forced to switch to flapping flight, or worse to land and take-off again, significantly increasing their energy expenditure [23,24]. While both thermals and carcasses are relatively unpredictable, thermals are not depletable contrary to carcasses. When a vulture discovers a carcass, its sharp drop in altitude while circling before landing is used as a signal by conspecifics, dragging tens of individuals to the food source in a few minutes [25,26]. As the number of vultures around the carcass increases (up to 100-120 individuals, [27,28]), individual feeding rates decrease due to reduced access to the resource, resource depletion by competitors and increased agonistic interactions [27]. Therefore, in these social birds, individuals should balance the advantage of conspecific presence to locate thermal updrafts [29] with the ultimate cost of competition around the carcasses that can be fierce [30–33]. As such, vultures are ideal models to investigate the role of conspecifics in shaping their foraging movement decisions.

Using a group of captive but freely-flying ‘griffon’ vultures, *Gyps fulvus* and *G. rueppellii,* tagged with high-resolution GPS loggers, we studied how conspecifics’ presence shapes individuals’ movement decisions during soaring flights. Despite being trained birds released for public shows, these individuals sometimes detected and fed on carcasses at surrounding farms (BN and YS, pers. obs.). We therefore consider these flights comparable to natural foraging flights. Focusing on the movement steps from thermal to thermal, we first assessed when do individuals preferentially discover new thermals (i.e. use of personal information) compared to using thermals already discovered by conspecifics (i.e. use of social information). We expected that vultures would favour the use of social information when unfavourable meteorological conditions increased thermal unpredictability and when flight conditions (e.g. low altitude) increased risks of landing [1]. Furthermore, given the hierarchy in vulture groups, we expected low-ranked individuals to be more prone to use personal information than high-ranked individuals to try to find the food source first, in order to avoid large aggregation [34,35]. Second, we investigated the drivers underlying thermal selection when individuals had to choose between simultaneously available thermals. We expected individuals to select thermals providing the maximal positive vertical speed (i.e. climb rate) as it may provide a reliable proxy of the thermal current strength helping them maximise their height gain [29]. To decrease uncertainty about resource finding and risks mentioned above, we expect that individuals should favour thermals hosting the maximum number of individuals to maintain cohesion and secure the possibility to cue on as many conspecifics as possible [29]. Finally, social preferences may also influence decision, with individuals preferentially moving together with preferred affiliates [12,36,37], as it could reduce competition due to familiarity between individuals [38].

## 2. Materials and methods

### 2.1 Study site, vultures housing conditions and experimental settings

The study was carried out in 2021 and 2022 at the Rocher des Aigles falconry centre, Rocamadour, France, and divided between winter and summer periods each year. During winters, vultures were housed within an aviary (6.7 x 6 x 6 m) equipped with four perches: three of them measuring 3.10 m, placed at 1.7, 2.6 and 3.5 m from the ground, and one of the full width of the aviary at 4 m height. This setting was used to estimate vulture social bonds (see Social bond estimation). In addition, besides being fed daily on small pieces of meat to prevent conflicts, five feeding events (one each week during a five-week period) were organised in the aviary on a butchery carcass occurring after a one-day fasting (to motivate feeding). These feeding events were used to assess dominance hierarchy within the group (see Hierarchy estimation). In summer, these trained vultures were kept perching on individual logs, released several times per day to execute free flight shows for the public within a landscape composed of plateaus interspaced by canyons, similar to “Causses” landscape typically used by french wild vultures [39]. The falconry centre is located near a 120 m-deep canyon and offers natural soaring conditions for raptors, making this study site a great place to investigate natural group flight behaviour (see Group flights), [24].

We used GPS data and visual observations to characterise the social and flight behaviour of eight captive vultures (7 Eurasian griffon vultures, *Gyps fulvus,* and 1 closely-related Rüppell’s vulture, *Gyps rueppellii*), including five females and three males (Table S1). Each year, we conducted experiments on a group of six individuals (two griffon vultures were replaced in 2022, Table S1). Experiments followed the animal ethic guidelines of France and the Centre National de la Recherche Scientifique. Handling of birds to fit GPS loggers followed the protocol of telemetry study of vultures authorised in the Programme Personnel 961, coordinated by OD, under the supervision of the French ringing centre, CRBPO, Paris. Furthermore, experiments, observations, handling and flight events were systematically performed under the guidance of the head of animal caretakers, BN.

#### 2.1.1 Social bond estimation

During five weeks in both years (December/January 2020-2021 and November/December 2021), we recorded pictures of vultures in the aviary from 8:00 to 19:00 (local time) at 5 minutes interval, using three camera traps (Wosport Big Eye D3 and Reconyx HyperFire HC600).

We identified birds using repeated colours on plastic rings and marks on the ruff and backhead feathers, using harmless colour sticks (Raidex GmbH, Figure 1A). We then processed recorded pictures to extract the individuals’ ID and position (bill or head position), using a purpose-built image annotation program in Julia software, JuliaHub Inc., [39]. For subsequent analyses, we relied on R software (v 4.2.2, R software, 2022, [40]).

**Figure 1.**
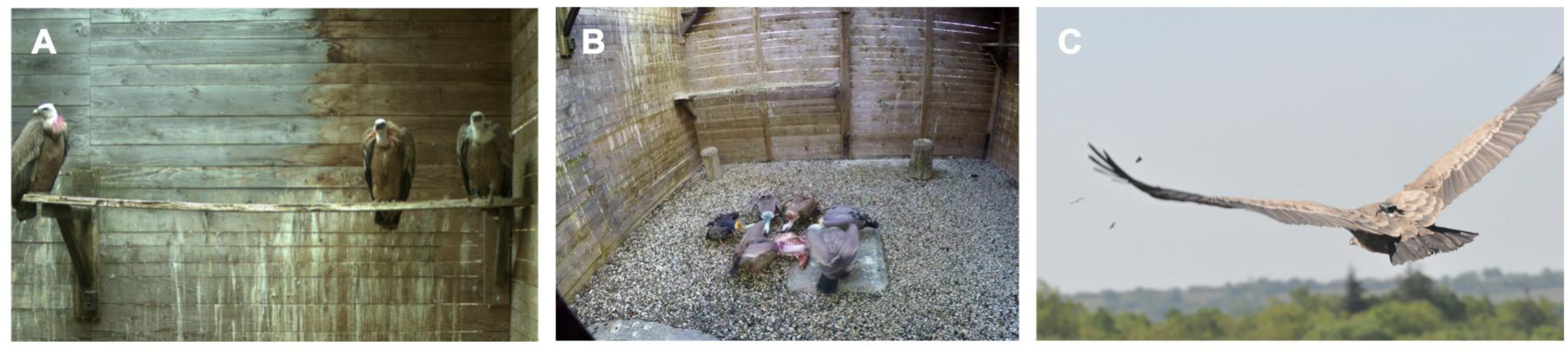
Data collection. (A) Perched vultures. Distance between vultures during perching events were used to estimate social-bond strength. (B) Feeding event around a butchery carcass. Agonistic interactions during those feeding events were used to estimate dominance hierarchy. (C) Flying vulture. Vultures were released for free flight into a 120-m canyon, equipped with high-resolution GPS loggers.

We considered the social bond between a dyad of individuals *i* and *j* based on spatial proximity following the Simple Ratio association Index (SRI, equation 1, [41,42])

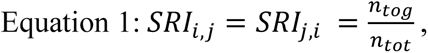

where *n_tog_* is the number of pictures in which individuals *i* and *j* were on the same perch at a Euclidean distance of less than 1.55 m and *n_tot_* is the total number of pictures in which individuals *i* and *j* were both detected on the same perch. SRI values varied between 0 and 1, where 0 represented dyads that were never seen associated and 1 represented dyads that were always observed sitting at less than 1.55 m from each other. The distance of 1.55 m was chosen as matching to the mode of the inter-individual distances distribution (Figure S1). This was also consistent with the aviary setting, as it corresponded to half the length of most available perches. Our analyses were robust to other choices for this distance threshold (see Supplementary Material, ESM01).

#### 2.1.2 Hierarchy estimation

Each winter, we estimated hierarchy within the vulture group by monitoring feeding interactions during the five carcass-based feeding events in the aviary (10 in total, Figure 1B) using a remotely-controlled video camera (GoPro Hero 4, GoPro Inc.) fixed at 2 m height on the aviary wall. These feeding events lasted on average 34 min (SD ± 4 min).

We computed individuals’ rank relying on the randomised Elo-rating approach [43,44], which accounts for potential temporal instability of the rank using permutations in the agonistic interaction series (‘elo_scores’ function, *aniDom* package, [44,45]; using 1000 randomisations and fixing the rank adjustment speed along the series, K-factor, to 200). The interaction series consisted in identifying the “wins” and “losses” for a given individual in agonistic interactions [46] with other individuals in each video (annotated with BORIS video analysis software, [47]). We used the ethograms from Bose & Sarrazin (2007, [30]) and Valverde (1959, [48]) to characterise griffon vulture feeding behaviour and between-individual interactions. An individual won the interaction when it interrupted another individual’s feeding bout (by pecking it, displacing it or engaging in a fight), and finally accessed the carcass before its opponent. In other cases, the interaction was considered as a “loss” for the initiator. We assessed the reliability of the dominance hierarchy through individual Elo-rating repeatability (‘estimate_uncertainty_by_repeatability’ function, *aniDom* package, [44]).

#### 2.1.3 Group flights

We recorded vulture flights decisions during 42 flight sessions (21 sessions each year) in the vicinity of the Rocher des Aigles. In general, birds were released for a flight session three times per day (in rare occasions from 2 to 4 times), at around 11:00, 14:30 and 16:00 (local time) for a mean duration of 26.03 min (SD ± 14.15 min) of flight. These captive vultures are trained to fly freely, searching for thermals, gaining altitude and coming back to their trainers (Supplementary Video 1). Vultures were equipped with a high-resolution GPS logger (4 Hz, TechnoSmart, models Gipsy 1, Gipsy 5 or Axytreck) positioned at their lower back using a Teflon leg-loop harness (Figure 1C, [49]). They were released in two groups, built according to social preferences with the three most socially-bonded birds together, at 2-min intervals. Release order alternated between consecutive days. For each flight session, we recorded and considered as stable the cloudiness (i.e. the proportion of clouds covering the sky, on a scale from 0 - no clouds - to 8 - sky fully covered by clouds), horizontal wind speed and temperature (all extracted from meteofrance.com based on local meteorological models).

To further investigate how vulture thermal choices were shaped by personal and social information, we pre-processed flight tracks in three consecutive steps. We subsampled individuals’ tracks from 4 to 1 GPS fix per second, and segmented their flight behaviour into gliding, linear soaring and circular soaring. We then created spatio-temporally dynamic maps of thermal availability based on the spatial clustering of individual’s circular soaring phases. Leaning on these maps, we retraced the history of thermal use/choice by individuals.

##### 2.1.3.1 Thermal use identification

To segment vulture flight between circular soaring, linear soaring and gliding flight we first calculated turning angle and vertical speed between consecutive locations using the *move* R package [50]. We applied a moving window of 30 s to calculate the absolute cumulative sum of the turning angles (hereafter cumulative turning angle) and a moving window of 5 s to calculate the average vertical speed. We then applied a k-means approach (k = 2, ‘kmeans’ function, *stats* R package) on the smoothed vertical speed (positive speed when flying upwards) to distinguish between soaring (ascending flight) and gliding (descending flight, [51,52]). We further classified soaring locations into circular soaring (indicating use of thermal updrafts) and linear soaring (also called slope soaring, expected to occur outside of thermals), with circular soaring being associated with a cumulative turning angle ≥ 300 degrees. A result of segmentation is illustrated in Figure 2B. Finally, we inferred the use of a thermal when the individual engaged in circular soaring for more than 30 s, with no interruption of more than 5 s of gliding.

**Figure 2.**
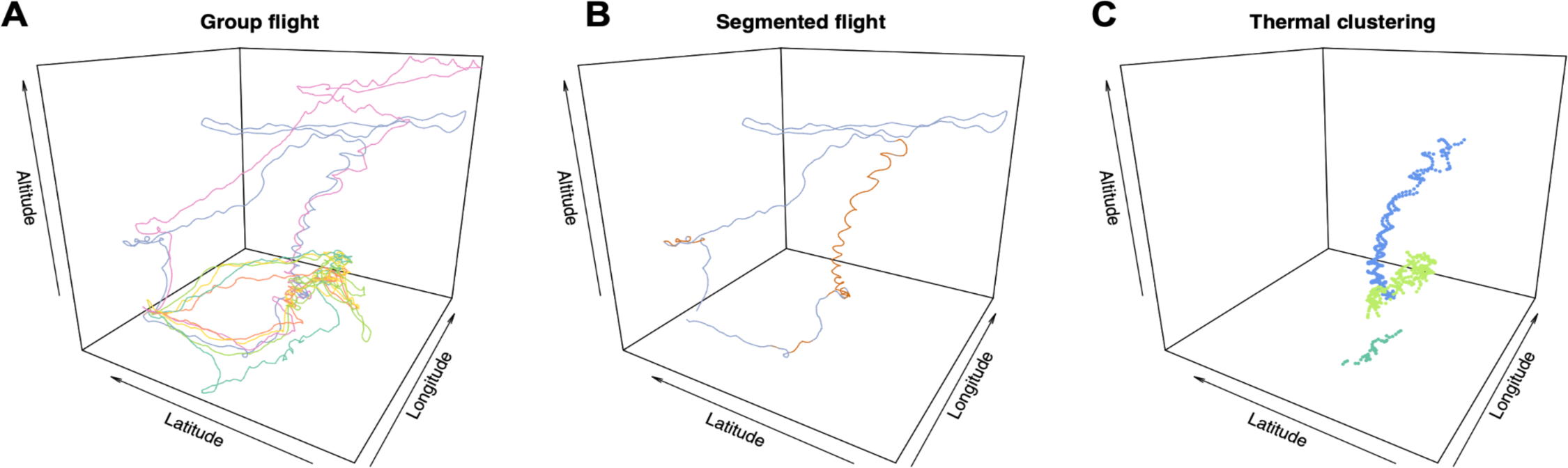
Flight data pre-processing. Pre-processing steps of group flight GPS data, example of one flight session. The altitude ranges from 200 m to 600 m. (A) shows a group flight (see Supplementary Video 1), with colours corresponding to each individual. (B) illustrates the segmentation of an individual’s flight (blue individual in (A)), with the orange segments corresponding to circular soaring phases. (C) illustrates the 3D density-based spatial clustering of individuals circular soaring phases, with colours indicating the three thermals identified in this flight session.

##### 2.1.3.2 Dynamic mapping of available thermals

Within each flight session, we created a dynamic map of thermals (Figure 3). First, we spatially clustered vulture circular soaring locations (reflecting the use of the same thermal updraft) independently of time by using a 3D density-based spatial clustering approach (’dbscan’ function, *dbscan* R package, [53]). This algorithm relies on a spherical neighbourhood to perform density-based neighbour joining, i.e. clustering (Figure 2C). We assumed this neighbourhood to be of a 40-m radius, and a minimum number of five locations within this range for the algorithm to consider the neighbourhood further. This 40 m threshold corresponded to the largest 4-nearest-neighbour distance observed when considering locations attributed to thermal use only (‘kNNdistplot’ function, *dbscan* R package) and matched with empirical expectations of radius during circular soaring phases [54].

**Figure 3.**
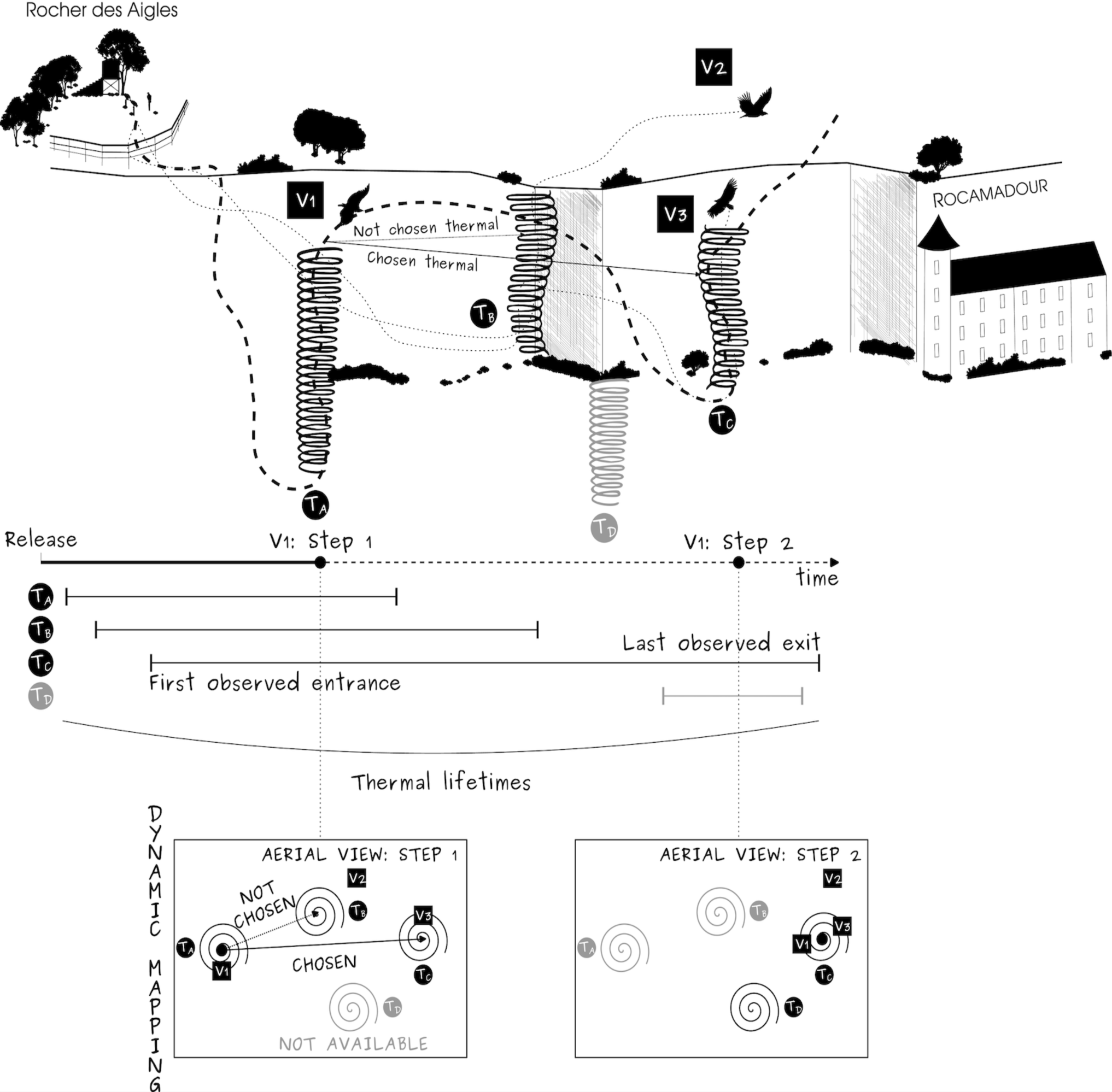
Illustration of the step selection framework used to investigate thermal selection. We focused on the movement of vultures released from the Rocher des Aigles when flying from thermal to thermal (i.e. a step). To do this, we mapped each thermal used during a flight session based on movement segmentation and clustering (see method section) to create dynamic maps of thermal availability over the flight session (as represented by the aerial views). The illustrated example focuses on the decision of a vulture (V_1_; step 1) when leaving the thermal (T_A_) and having to choose between two available thermals (T_B,_ close but not currently used by another vulture, and T_C_, further away but currently used by another individual). T_D_ was not available until step 2, when it was discovered and used by another individual, and is therefore shown in grey at step 1. At step 2, V_1_ joined V_3_ in T_C_ and both thermals T_A_ and T_B_ were no longer available. A thermal was available from the moment when the first individual entered it until the last individual left it. Therefore, the number of available thermals could change during the flight session (see differences between maps in step 1 and 2).

We then made those maps dynamic in time by considering the lifetime of each thermal. We considered a thermal as “available” from the moment when the first individual entered it until the last individual left it (Figure 3).

### 2.2 Statistical analyses

We defined collective flight events as any time of a flight session when at least two individuals were flying. For each of these events, we first analysed the use of social information (the tendency to join thermals already discovered by conspecifics) as a function of external (meteorological) and internal drivers (individual traits). We then used step selection functions to define, at each movement step, which drivers determined the selection of the chosen next location (thermal updraft) relative to other potential locations.

#### 2.2.1 Drivers of social information use

We investigated the effect of local meteorological context, individual traits and flight mechanics on the use of social information, defined here as the tendency to join thermals already discovered by conspecifics. We considered that an individual discovered a thermal when it was the first, among all individuals, to adopt circular soaring flight into it. For the analysis, we discarded the discovery of the first thermal in each flight session (as this thermal was necessarily discovered).

To investigate the drivers underlying the use of social information we modelled the probability to join a thermal already discovered by others using generalised linear mixed models (GLMMs) with binomial error structure and a logit link function [55]. Our full model contained the following ten fixed effects: **meteorological variables** with the (i) wind speed (categorical predictor), (ii) cloudiness and (iii) temperature (both continuous predictors); **social variables** with (iv) the age (continuous predictor) and (v) rank in the dominance hierarchy of the individual (ordinal categorical predictor), and variables related to the **mechanic of flight** with (vi) the glide-ratio (horizontal distance travelled during a 1-m altitude loss, only measured on glides with straightness > 0.95 in each flight), (vii) the altitude of and (iix) the 3D distance to the exit location from the previous thermal used (all continuous predictors). We also added (ix) the group in which individuals have been released (first or second group released for the flight) and (x) the time elapsed since the first individual take-off (continuous predictor) as control variables. Individual ID was considered as a random factor.

To compare the relative importance of the fixed effects we scaled all non-categorical variables to use their estimate as dimensionless effect size [56]. We examined the significance of each variable by comparing the goodness of fit of models with and without the variable of interest using a likelihood ratio test (‘drop1’ function, *stats* R package). Assumptions required for these statistical approaches (homoscedasticity, Gaussian distribution of residuals) were checked with plot diagnosis (histogram of residuals, residual Q-Q plot, distribution of residuals vs fitted values, *DHARMa* R package, [57]). We also tested for the presence of outliers, and calculated the variance inflation factor (VIF) to test for collinearity (VIF values ≥ 3 suggesting a strong collinearity [58]). We did not detect collinearity in our predictors (VIF^max^ = 1.74) (Figure S3). Furthermore, we extracted the marginal coefficient of determination (R_m_^2^) and the conditional coefficient of determination (R_c_^2^) which describe, respectively, the proportion of variance explained by fixed effects and by the fixed and random effects combined [59]. Finally, as the flight time period, and the tested individuals differed, we cross-compared models fitting on the two years separately (see Supplementary Material ESM01).

#### 2.2.2 Drivers of thermal updraft selection

To study the drivers underlying thermal selection, we embedded our work in the Step Selection framework [60]. We considered the series of thermals used by each individual. In that series, we focused on movement steps involving a flight to a thermal previously (or currently) used by a conspecific when other thermals were available. Using a conditional logistic regression, we compared the “chosen” thermal characteristics to those “available” but not chosen. The conditional logistic regression included seven predictors, respectively characterising the **thermal profitability** with (i) the distance to it and (ii) maximum vertical speed reached in the thermals by any individual since the focal individual has been released in the flight session (continuous predictors), **individual personal experience** considering whether (iii) the thermal was previously used by the focal individual (binary predictor), and **social information** with (iv) the presence of the focal individual’s preferred affiliates in the thermal or not (binary predictor), (v) the number of individuals present in the thermal, (vi) the weighted mean (by the number of previous visits to the thermal) of the social bond with individuals that used the thermals, and (vii) the negative cubed difference of ranks between the focal individual and those in the thermals (all continuous predictors, set to 0 for the two latter if no individuals used it/were present). We used the negative cubed difference to consider an attraction-repulsion effect (high rank toward low rank and the opposite respectively) by translating a linear rank difference to a relative hierarchy scale which enhances large rank differences. For example, following the curve of the negative cube function, if the difference of rank was five (e.g. the focal individual is ranked 6^th^ - a low rank, a conspecific in another thermal is ranked 1^st^ - a high rank) the probability that the focal individual joined the conspecific should be drastically decreased, mimicking a repulsion effect.

Also for this model, we scaled all non-categorical variables to better compare their relative importance. We fitted the conditional regression considering all individuals together, yet considering data stratified at the individual-step level. We finally reported the relative selection strength (RSS) of significant variables which provides the magnitude of estimated selection coefficients, holding all other covariates fixed [61,62].

## 3. Results

Vulture dominance hierarchy was steep (Figure S2) and reliably inferred (individual Elo-rating repeatability = 0.82 and 0.83 in 2021 and 2022 respectively). The rank orders among individuals present in both years were relatively consistent and unrelated to sex or age (Table S1). During the 21 flight sessions performed each year, we identified a total of 520 and 578 thermalling events in 2021 and 2022. On average, 63% (SD ± 7%, Table S1) of these circular soaring behaviours took place in thermals discovered by a conspecific.

### 3.1 Flight risks and hierarchy shapes the use of social information

Our model was significantly better than the null model (considering only control effects; AIC = 1237.4 and 1412.7 respectively) and explained 30% of the variance (Table S2). The probability for an individual to use a thermal previously discovered by a conspecific decreased with temperature (from 0.74 at 17°C to 0.43 at 31°C, Figure 4A, Figure 5A, Table S2), but tended to increase with cloudiness and wind speed (Figure 4A, Table S2). This probability dropped also with the distance from the previous thermal and the altitude at which the bird left it (from 0.63 when being at a distance of 12 m from the last thermal used to 0.16 at a distance of 6776 m and from 0.76 when exiting the last thermal at an altitude of 195 m to 0.039 at 1574 m of altitude, Figure 4A, Figure 5B, C, Table S2). Individuals lower in the dominance hierarchy were approximately twice as likely to discover new thermals than high-ranked individuals (Figure 4A, Figure 5D). We did not detect significant effects of age and glide-ratio on the probability to use thermal previously discovered by conspecifics (Figure 4A, Table S2). Fitting the same model structure on 2021 and 2022 data separately yielded the same overall results, suggesting that the observed pattern was robust to changes in hierarchy and between-year conditions (Figure S4, Table S2).

**Figure 4.**
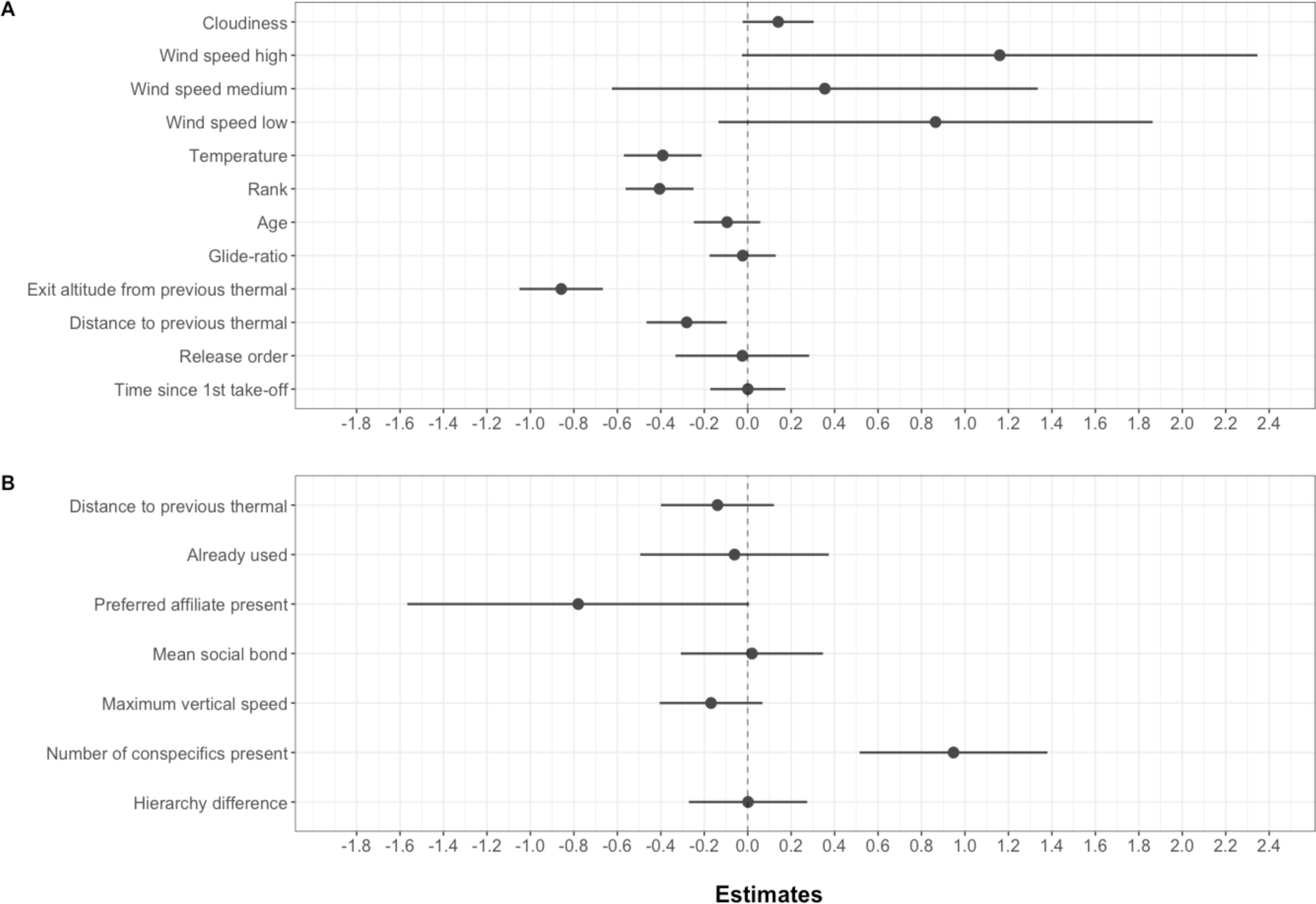
Estimates of models investigating the drivers of social information use (A) and thermal selection (B). Rows correspond to each predictor. Each point represents the standardised estimate value. Segments give the associated 95% confidence intervals.

**Figure 5.**
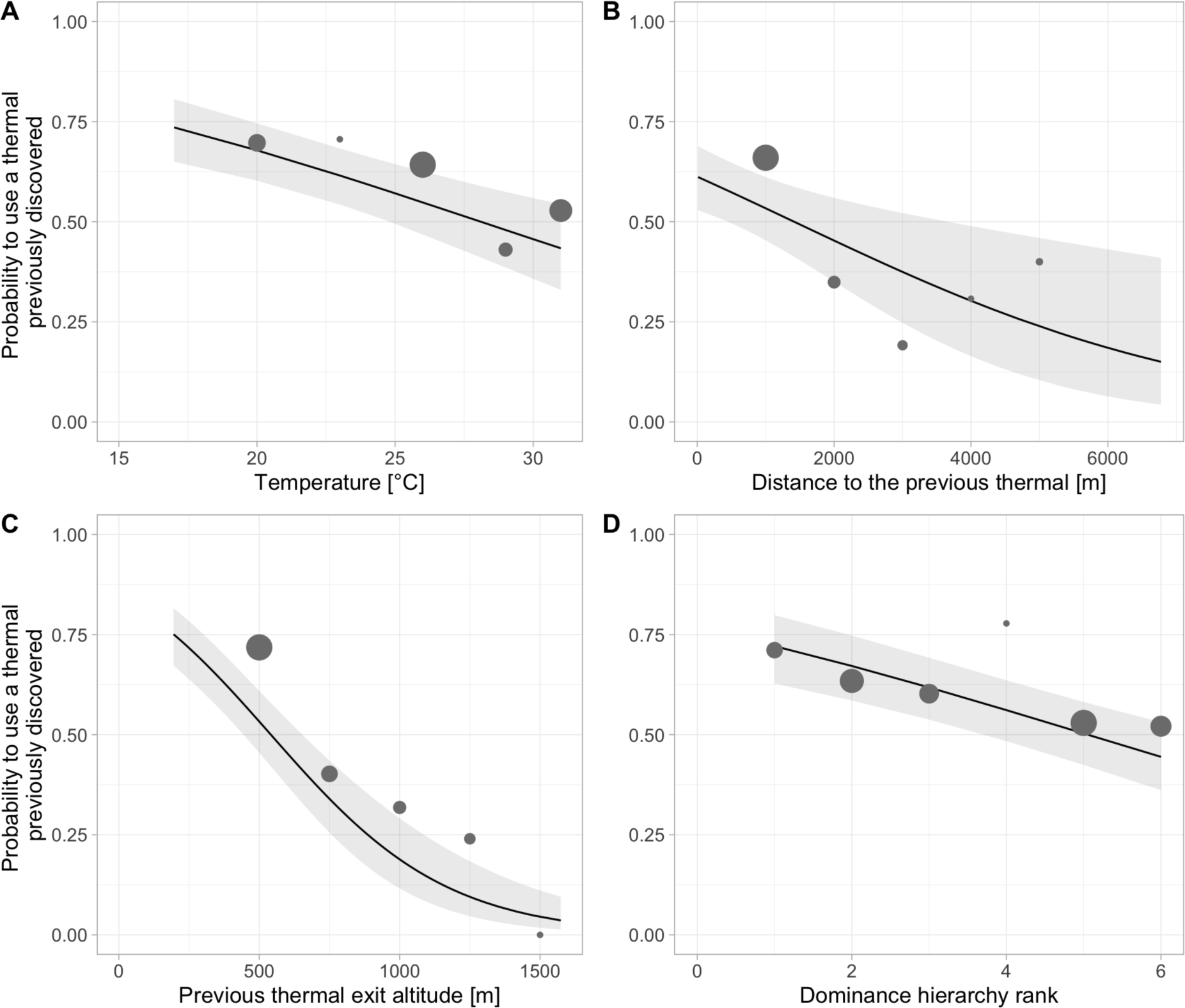
The probability to use a thermal already discovered by conspecifics decreases with temperature, distance to the thermal, flight altitude and hierarchy rank. Points represent the probability of using a thermal already discovered by a conspecific, estimated on the raw data. Their size is relative to the sample size. To do so, for (A), (B) and (C), predictors were binned. Black lines with grey shades show the GLMM estimated probability with its 95% confidence interval (N = 1098 thermals).

### 3.2 Vultures select thermal updrafts hosting the most conspecifics

We identified 178 movement steps where an individual entered a thermal while at least one other thermal was available simultaneously. Individuals were significantly more likely to select thermals hosting the largest number of conspecifics at time of decision (Relative Selection Strength [95% confidence interval] = 27.94 [5.99, 131.63]; Figure 4B, Table S3). On the contrary, the probability to choose a thermal tended to decrease when the preferred affiliate was using it. The distance to the previous thermal and the maximal vertical speed reached in the thermal and whether individuals used this thermal in the past did not significantly affect thermal selection (Figure 4B, Table S3). At time of decision, the difference in dominance ranks as well as the presence of its preferred affiliate did not drive the individual’s probability of selecting the thermal. This pattern was consistent when considering only movement steps where individuals had to choose between thermals used at time of decision (N = 61, Figure S5, Table S3). Furthermore, considering all decision events, the sensitivity analysis on the inter-individual distance threshold for the social bond strength estimation yielded the same results (i.e. 1.55 m, 1.30 m and 1 m; see Supplementary Material ESM01, Figure S6, Table S4).

## 4. Discussion

Using a combination of high-resolution tracking and social structure monitoring, we identified contextual drivers for the differential weighting of personal and social information in movement decisions. We showed that vultures’ movement decisions predominantly relied on social information, especially in unfavourable flight conditions that increased thermal unpredictability or put individuals at risk of undesired landing. Overall, individuals preferentially joined thermals with the largest number of conspecifics. However, the use of social information depended on the individual social status: low-ranking individuals were more inclined to use personal information and discovered more thermals on their own than high-ranking individuals.

We found that low-ranked individuals, likely the ones suffering the most from interference competition, had higher probabilities of discovering new thermals, thus likely exploring their environment more intensively than the high-ranked individuals. Such flight strategy would enable subdominant individuals to reach carcasses first, or at least to arrive at the beginning of the feeding event when the rate of interference is lower [27] hence avoiding to loose opportunities due to conformity with conspecific behaviour [64]. From this may emerge a producer-scrounger dynamic [65,66] wherein the use of personal information from low-ranked individuals to arrive at food sources with lower competition levels would be exploited by dominant individuals to reduce their own searching effort [16,66,67]. This is coherent with previous observations of low-ranked vultures being “pioneers”: the very first individuals to land and feed on the carcasses before being displaced by high-ranked individuals arriving afterwards [27]. This influence of dominance on foraging tactics where low-ranked individuals explore and find food while dominant profit has also been observed in other social bird species such as common cranes *grus grus*, oystercatcher *Haematopus ostralegus*, house sparrows, *Passer domesticus*, and barnacle goose, *Branta leucopsis*, [18,67–69]. Eviction of subordinates from food patches has even recently been identified as a trigger for collective movements [70]. In contrast, in activities where individuals do not experience competition, such as tool-use learning in chimpanzees, naïve individuals will generally copy dominant (and knowledgeable) individuals [71]. Our study hence stands as a clear cut illustration of the “copy when asocial learning is costly’’ rule [72]: the vulture position in the dominance hierarchy, through the costs it imposes on access to food, seems to calibrate the balance between the use of personal and social information in foraging movements. In some cases, however, trading personal information in favour of social information is inevitable.

When the environment is largely unpredictable or whenever using error-prone personal knowledge can be energetically costly, individuals should tend to eavesdrop, and rely more on information provided by conspecifics to reduce uncertainty about resources availability [15,73]. Here, we evidenced both cases. First, vultures prioritised the use of social information when the temperature was low and tended to when cloudiness and wind speed increased. These weather conditions may translate into fewer and weaker thermals, drifting into the wind, making them less predictable [74–78]. Second, they also favoured social information when the altitude at which they left their previous thermal was low. When exiting a thermal at low altitude, individuals have limited time to glide to the next thermal before having to shift to flapping flight to stay aloft, or else landing in an undesired place, which both would add high energetic cost associated with flapping and take-off [23,24,79,80]. Reaching high altitudes quickly to avoid this risk may also explain why vultures used more thermals previously discovered by conspecifics if those were close to the last thermal they used. While vultures are able to cope with difficult flight conditions (e.g. turbulence and strong wind) by adjusting their banking angles [55], anticipating such risky events may remain the most efficient way to maximise the trade-off between time, energy and risk which largely dictates their flight strategy [81]. Adult individuals, through experience, are generally better at coping with difficult flight conditions [82], yet we did not evidence an effect of age relative to the use of social information, as observed in other group living species (e.g. [83]). More than age *per se,* the familiarity of individuals with a given situation might shape their tendency to rely or not on social knowledge (e.g. in spider monkeys, *Ateles geoffroyi*, during collective foraging [84]). The captive individuals tested in this experiment are all adults and fly in the same landscape every day since their birth, thus they are probably very familiar with the areas favourable to thermal emergence. This could explain why we did not detect any effect of age on the use of social information, but also indicates that the relative importance of this source of information is probably underestimated due to the birds’ familiarity with the surroundings. An interesting complementary experiment (though technically difficult) to disentangle the effect of familiarity with the landscape would be to move the whole group and repeat the experiment in an unfamiliar area to better estimate the strength of personal versus social information use.

When faced with a choice between simultaneously available thermals, the previous experience of individuals (i.e. whether the thermal was used previously or not by the focal) or current expertise of the group (i.e., relative age/hierarchy difference) impacted very little vulture movement decisions compared to other social cues, contrasting with previous findings from insects to mammals, including birds [85–89]. In the current system, ascending currents can be very ephemeral phenomena, sometimes only lasting a few minutes [90,91]. Certainly, a “live report” is therefore better provided by the accumulation of convergent information sources (i.e. numerous conspecifics, [92]) rather than relying on a unique individual source (i.e. the individual itself or one reference individual). In that line, and surprisingly, the presence of one preferred affiliate in a thermal tended to reduce the probability to join it. There is evidence that social bonds assessed “on the ground” are often unrelated to co-flight preferences [93]. It therefore questions whether collective flights might be used by vultures to strengthen initially weak social bonds. Maintaining in-flight bounds can indeed be important, as evidenced in the migratory behaviour of other soaring bird species to enable accurate collective mapping of the distribution of uplifts [94,95]. Furthermore, for soaring birds, the presence of conspecifics should provide not only information on the location and strength of updrafts [20,95] but could also indicate flight speed and circling radius needed to optimise climb rate, by remaining close to the centre of the thermal where uplift is highest [55]. Yet, the maximum speed reached by individuals using the thermal little affected vulture decision choices. Possibly, climb rate or individual speed are not as easy to assess at a distance, compared to the number of conspecifics. In other words, vultures tended to favour quantity signals (with the number of conspecifics) over quality signals (maximal vertical speed) [96]. The “power of the group” may indeed in turn drive cohesion, which could itself make social information even more profitable [96,97].

Altogether, our results provide insights into the architecture of decision-making during movement in a social bird. It highlighted the trade-offs between personal and social information these birds have to consider in order to optimise both their flying efficiency and their foraging success. As a first approximation, we considered social cues as coming from “conspecifics”. Strictly speaking however, our study included two species, Griffon vulture and Rüppell’s vulture, albeit phylogenetically close and with similar biology. The one Rüppell’s vulture in fact, used social information provided by surrounding vultures and did not stand out as an outlier in its behaviour. It is known that even phylogenetically distant individuals could be an important source of social information, not only about the presence of carcasses [98], but also about the availability of thermals when sharing the same airspace (e.g. from black kites, *Milvus migrans,* or common swifts, *Apus apus,* [99,100]). Interactions with heterospecifics can indeed drastically affect animals’ daily life [101], up to shaping the cognitive machinery underpinning their foraging decisions [102]. How heterospecific cues are used when foraging remains clearly overlooked. Future studies in this direction could provide valuable insights into understanding the fundamental rules dictating how animals decide where to go.

## CRediT authors’ contributions

**Yohan Sassi**: Conceptualization, Methodology, Software, Investigation, Formal Analysis, Visualization, Writing - original draft, Writing - Review & Editing.

**Basile Nousières**: Investigation, Resources.

**Martina Scacco**: Software, Writing - Review & Editing.

**Yann Tremblay**: Conceptualization, Writing - Review & Editing.

**Olivier Duriez**: Conceptualization, Investigation, Supervision, Writing - Review & Editing.

**Benjamin Robira**: Conceptualization, Methodology, Software, Visualization, Validation, Supervision, Writing - original draft, Writing - Review & Editing

## Data availability

Scripts for review and supplementary video are available here: https://github.com/YohanSassi/UpdraftsDecisions

A perennial storage of data and scripts will be provided after revision (e.g. Zenodo, Dryad)

## Acknowledgment

We thank Dominique Maylin and Raphael Arnaud at Rocher des Aigles as well as all of the staff for their patience, help and enthusiasm for the project. GPS loggers were provided by Giacomo Dell’Omo. Camera traps and Gopro video cameras were provided by Aurélien Besnard, Samuel Caro and Samuel Perret. We also thank Samson Acoca-Pidolle for fruitful discussions about the statistical analyses.

