## Supplementary materials for "The use of social information in vulture flight decisions"

a: CEFE, Univ Montpellier, CNRS, EPHE, IRD, Montpellier, France
b: Rocher des Aigles, Rocamadour, France
c: Max Planck Institute of Animal Behavior, Department of Migration, Am Obstberg 1, 78315, Radolfzell, Germany
d: Department of Biology, University of Konstanz, Universitätsstraße 10, 78464, Konstanz, Germany
e: Marine Biodiversity Exploitation and Conservation (MARBEC), University of Montpellier, CNRS, Ifremer, IRD, 34203 Sète, France
f: Department of Biodiversity and Molecular Ecology, Research and Innovation Centre, Fondazione Edmund Mach, San Michele all'Adige, Italy

§ Co-last authors

### Supplementary materials

#### ESM01: Sensitivity analysis

To assess the robustness of our inference to variability in the social composition of the group and flight conditions between years and other “arbitrary” analytical decisions, we repeated the analysis based on different choices.

Drivers of social information used in movement decisions were investigated on all data combined to increase sample size. Yet, we evaluated the effect of years by refitting the full model structure on 2021 and 2022 data separately.

To study the drivers of thermal selection, we investigated the influence of social bond strength between individuals on thermal choice. Social bond strength was calculated considering spatial proximity on perches (≤ 1.55 m) between individuals as a sign for social bond. We repeated the model fit (see main text, Drivers of thermal updraft selection) using a distance threshold of 1.30 m (half the average wingspan of griffon vultures, [1]), and 1 m.

Furthermore, in the main text, to analyse thermal updraft choices, we considered all choices independently of whether vultures had to choose between thermals with and potentially without conspecifics. To verify whether the “presence” only of conspecifics influenced the choice, we repeated the analysis considering only choice events implying that currently available thermal updrafts were all used by conspecifics.

All these investigations revealed high robustness of our inference (Figure S5-7, Table S2-4). Therefore, we presented in main text the models which were the most complete and based on the most parsimonious rationale.

**Supplementary Video 1** is accessible with scripts at:
https://github.com/YohanSassi/UpdraftsDecisions

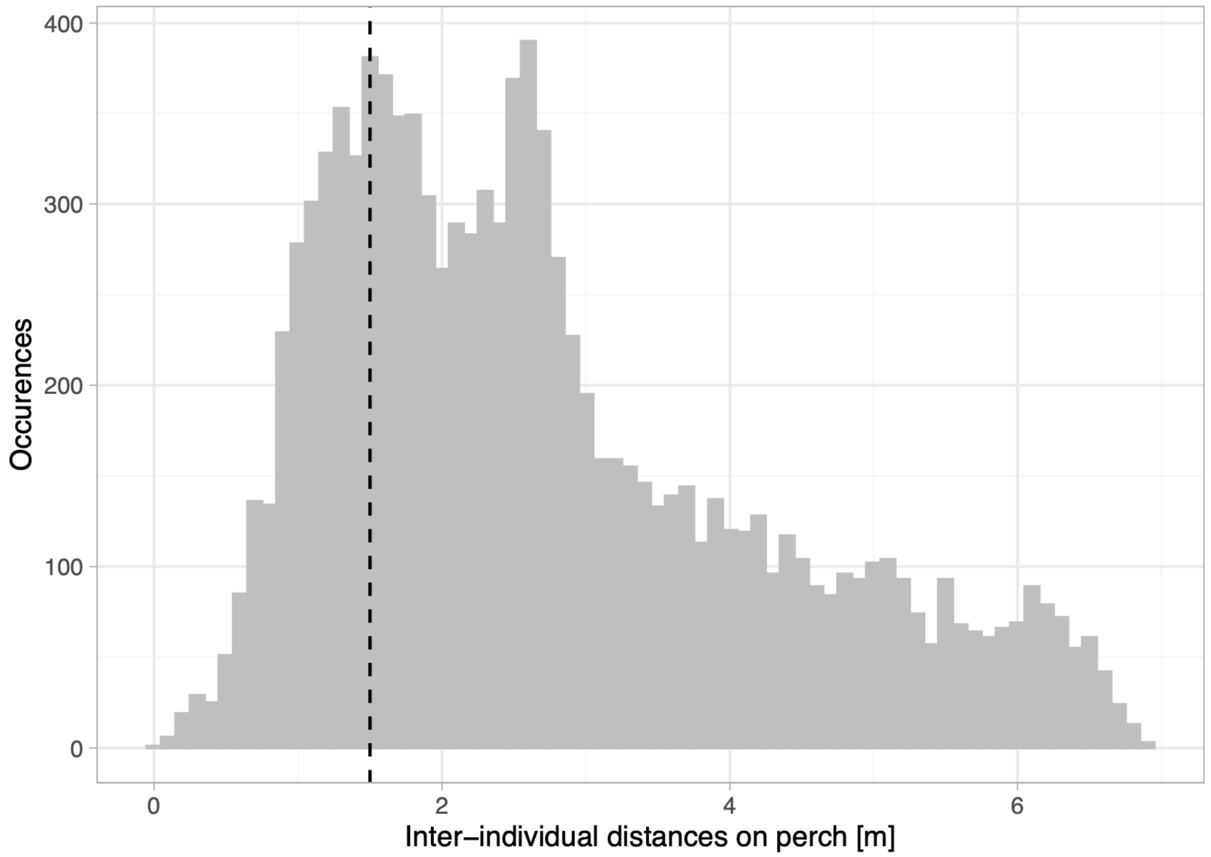


**Figure S1. Distribution of the Euclidean inter-individual distances between birds on perches.** Distances have been measured based on pictures taken in the wintering aviary. Every bin corresponds to 10 cm. The vertical black dotted line represents a distance of 1.55 m.

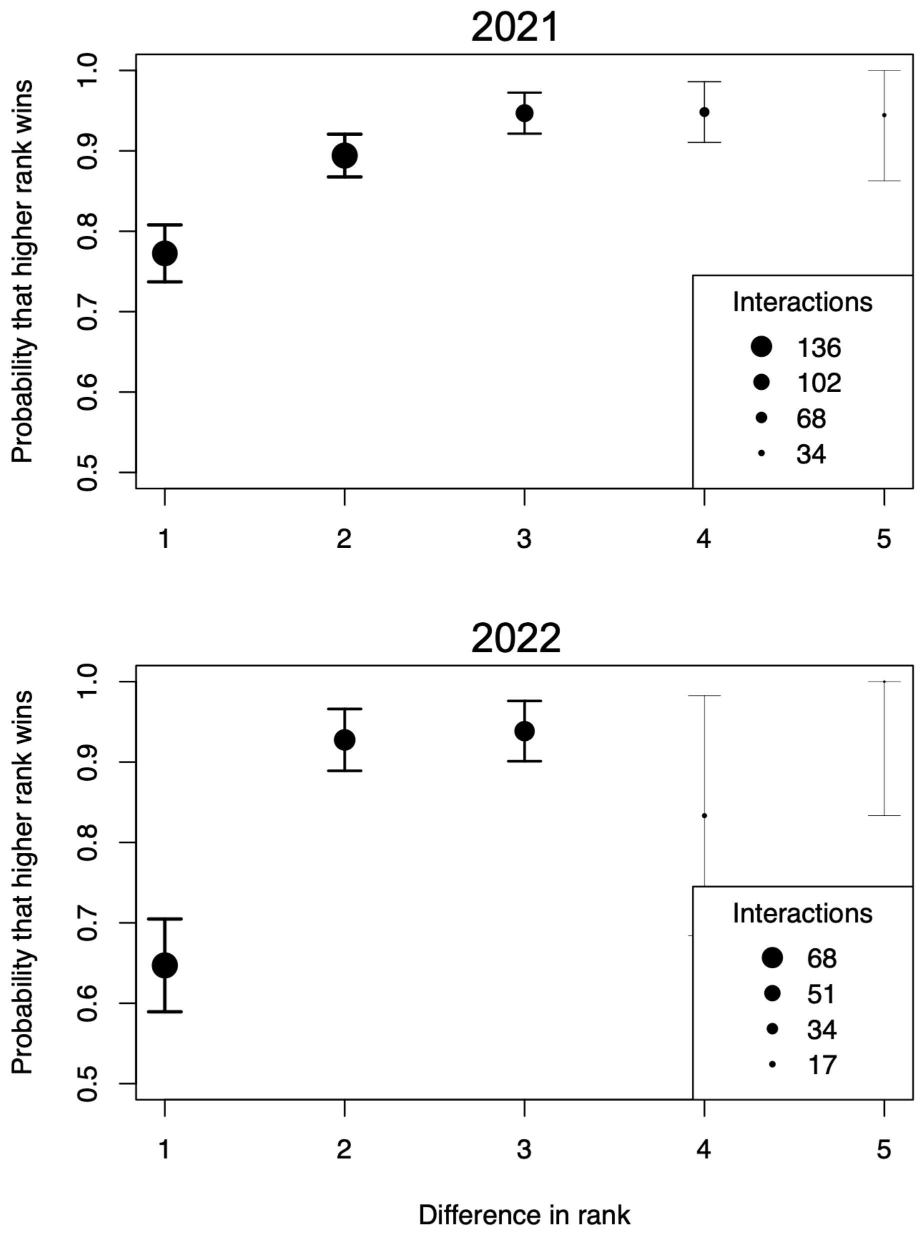


**Figure S2. Dominance hierarchy steepness in vulture groups.** Points represent the probability that the higher ranked individual wins an agonistic interaction as a function of the difference in rank. The size of the point is relative to the sample size. Segments give the 95% confidence interval.

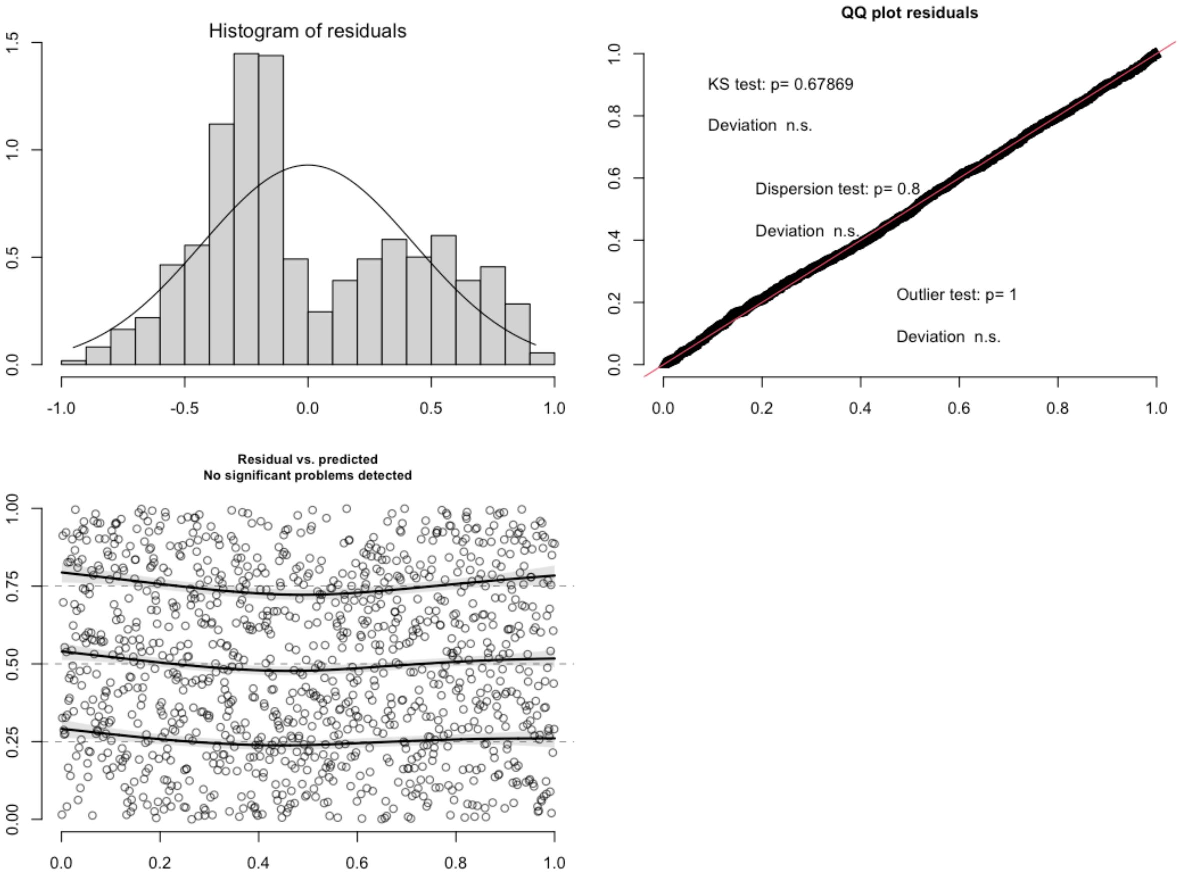


**Figure S3. Diagnostic plot of the model investigating the drivers of social information use.**

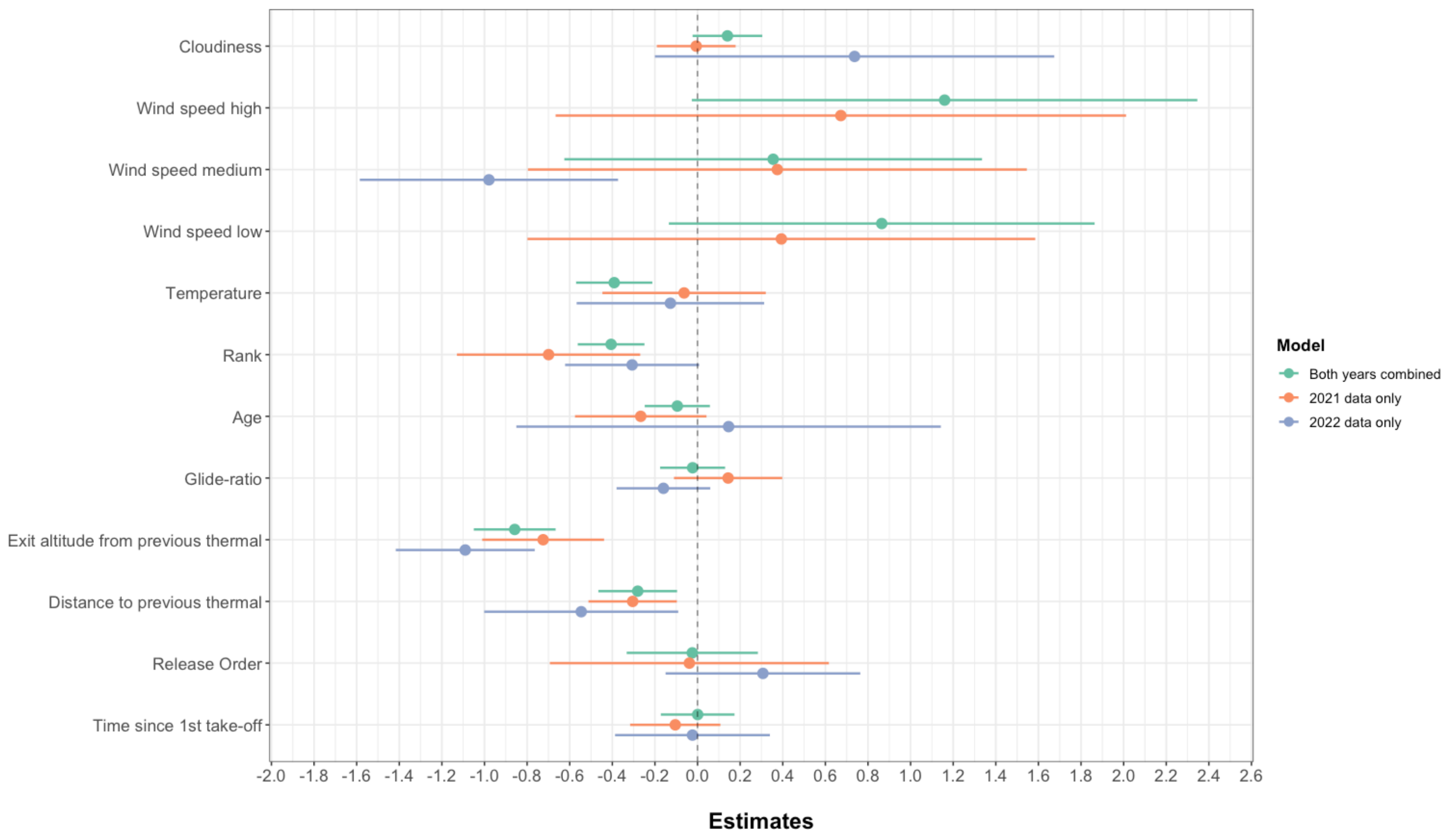


**Figure S4. Estimates of models investigating the drivers of thermal discovery.** Rows correspond to each predictor. Each point represents the (standardised) estimate value. Segments give the associated 95% confidence intervals. Models were fitted considering data of both years (green), only 2021 (orange) or only 2022 (blue).

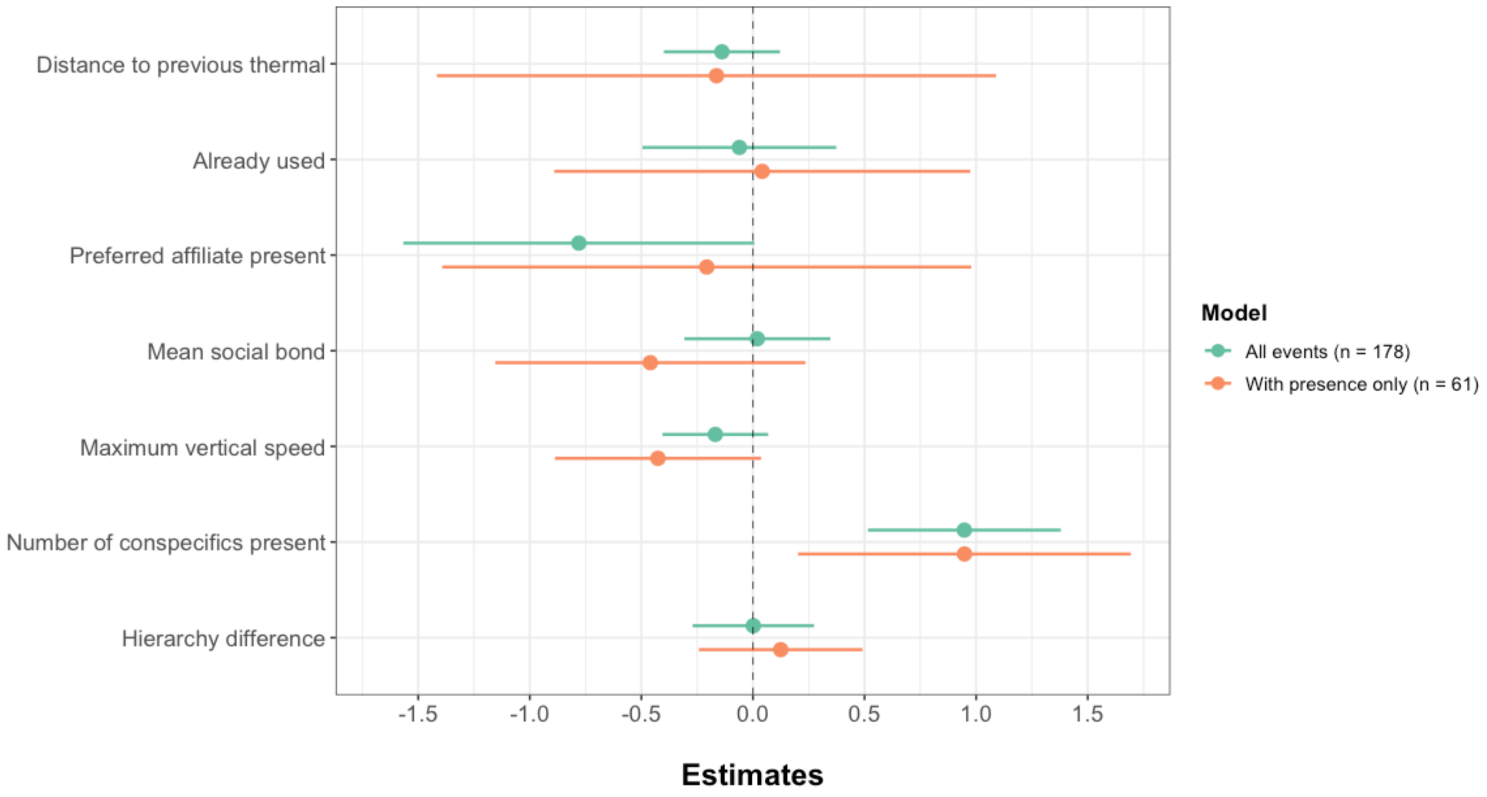


**Figure S5. Estimates of models investigating the drivers of thermal selection.** Rows correspond to each predictor. Each point represents the (standardised) estimate value. Segments give the associated 95% confidence intervals. “All events" refers to the model considering decision events where available thermals previously used but potentially empty at time of choice were considered. “With presence only” refers to the model focusing on decision events where vultures had to choose only between thermals currently used by conspecifics.

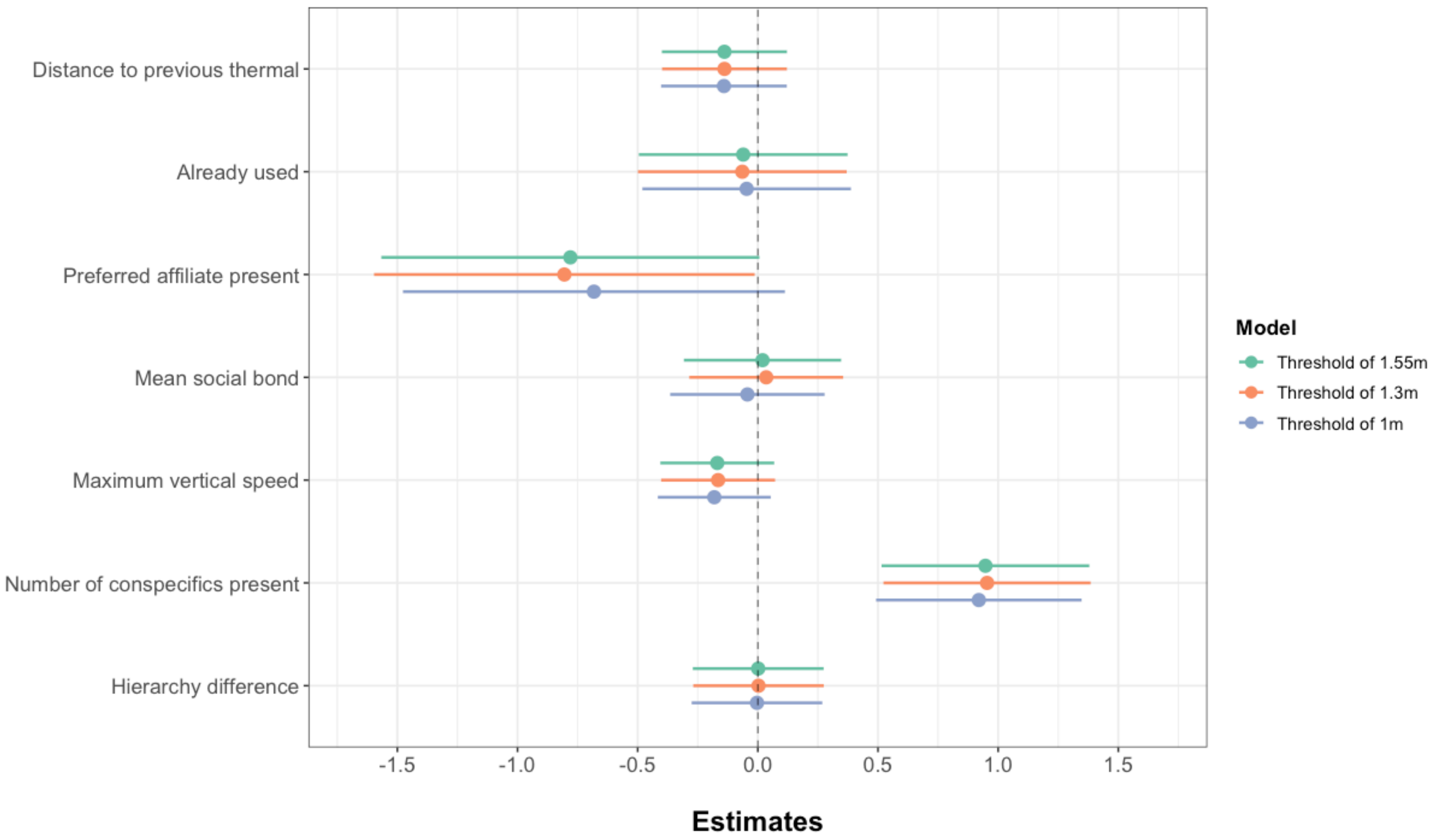


**Figure S6. Estimates of models investigating the drivers of thermal selection.** Rows correspond to each predictor. Each point represents the (standardised) estimate value. Segments give the associated 95% confidence intervals. "Threshold 1.55m" refers to the model considering an inter-individual distance threshold of 1.55m in the estimation of social bonds, which is the threshold considered in the analysis. The other levels are part of the sensitivity analysis.

**Table S1. Summary table of vultures under study**. For ranks, (-) means that for the considered year the rank of the individual was not estimated. The frequency of socially informed thermal use corresponds to the percentage of thermals used by each individual that was previously discovered by a conspecific.

| Individuals | Species | Sex | Age | Year of presence | Rank (in 2021 / 2022) | Frequency of socially informed thermal use | Number of thermalling events |
|---|---|---|---|---|---|---|---|
| **Henry** | *G. fulvus* | F | 25 | 2021 | 3 / - | 0.654 | 78 |
| **Gregoire** | *G. fulvus* | M | 11 | 2021 | 1 / - | 0.716 | 81 |
| **Hercule** | *G. fulvus* | F | 10 | 2022 | - / 4 | 0.667 | 27 |
| **Kazimir** | *G. fulvus* | M | 8 | 2021 / 2022 | 6 / 6 | 0.521 | 190 |
| **Bulma** | *G. fulvus* | F | 8 | 2021 / 2022 | 4 / 3 | 0.652 | 138 |
| **Kirikou** | *G. rueppelli* | F | 8 | 2021 / 2022 | 5 / 5 | 0.529 | 291 |
| **Leon** | *G. fulvus* | M | 6 | 2021 / 2022 | 2 / 2 | 0.634 | 246 |
| **Mathilda** | *G. fulvus* | F | 4 | 2022 | - / 1 | 0.702 | 47 |

**Table S2. Outputs of models investigating the drivers of social information use.** Estimates for continuous variables are scaled and provided with their 95% confidence intervals (95% CI), hence can be used as a dimensionless effect size. For the categorical predictor wind speed, the level without wind (i.e. wind speed null) is included in the intercept. ID gives the percentage of variance explained by the individual names (random effect), with $N_{ID}$ the number of individuals considered. (-) indicates missing data preventing estimation of the considered parameters. Scaled variables are indicated by an asterisk.

| | **Use of already discovered thermal** | | | | | | | | |
|---|---|---|---|---|---|---|---|---|---|
| | *Both year combined* | | | *2021 data* | | | *2022 data* | | |
| *Predictors* | *Estimates* | *95% CI* | *p* | *Estimates* | *95% CI* | *p* | *Estimates* | *95% CI* | *p* |
| (Intercept) | -0.24 | [-1.18, 0.71] | - | 0.61 | [-0.54, 1.77] | - | 0.52 | [-0.40, 1.44] | - |
| Cloudiness * | 0.14 | [-0.02, 0.30] | 0.089 | -0.01 | [-0.19, 0.18] | 0.947 | 0.74 | [-0.20, 1.67] | 0.120 |
| Wind speed high | 1.16 | [-0.03, 2.35] | | 0.67 | [-0.67, 2.01] | | - | - | |
| Wind speed medium | 0.35 | [-0.63, 1.34] | **0.011** | 0.37 | [-0.80, 1.55] | 0.792 | -0.98 | [-1.59, -0.37] | **0.001** |
| Wind speed low | 0.86 | [-0.13, 1.86] | | 0.39 | [-0.80, 1.59] | | - | - | |
| Temperature * | -0.39 | [-0.57, -0.21] | **<0.001** | -0.06 | [-0.45, 0.32] | 0.747 | -0.13 | [-0.57, 0.31] | 0.570 |
| Dominance rank * | -0.41 | [-0.56, -0.25] | **0.003** | -0.70 | [-1.13, -0.27] | **0.017** | -0.31 | [-0.62, 0.01] | 0.054 |
| Age * | -0.09 | [-0.25, 0.06] | 0.247 | -0.27 | [-0.58, 0.04] | 0.134 | 0.15 | [-0.85, 1.14] | 0.774 |
| Glide-ratio * | -0.02 | [-0.18, 0.13] | 0.767 | 0.14 | [-0.11, 0.40] | 0.266 | -0.16 | [-0.38, 0.06] | 0.158 |

| | | | | | | | | | |
|---|---|---|---|---|---|---|---|---|---|
| Exit altitude from previous thermal * | -0.86 | [-1.06, -0.66] | **<0.001** | -0.72 | [-1.01, -0.44] | **<0.001** | -1.09 | [-1.42, -0.76] | **<0.001** |
| Distance to previous thermal * | -0.28 | [-0.47, -0.10] | **0.002** | -0.30 | [-0.51, -0.10] | **0.003** | -0.55 | [-1.00, -0.09] | **0.012** |
| Release order [2] | -0.02 | [-0.33, 0.28] | 0.875 | -0.04 | [-0.69, 0.62] | 0.908 | 0.31 | [-0.15, 0.76] | 0.186 |
| Time since 1st take-off | 0.00 | [0.17, 0.17] | 0.994 | -0.10 | [-0.32, 0.11] | 0.339 | -0.02 | [-0.39, 0.34] | 0.898 |
| *Random Effects* | | | | | | | | | |
| ID | 0.00 | | | 0.18 | | | 0.00 | | |
| $N_{ID}$ | 8 | | | 6 | | | 6 | | |
| Observations | 1098 | | | 520 | | | 578 | | |
| Marginal $R^2$ / Conditional $R^2$ | 0.304 / - | | | 0.269 / 0.308 | | | 0.406 / - | | |
| ** Mean + SD prior to scaling* | | | | | | | | | |
| Cloudiness | 1.07 + 1.73 | | | 1.98 + 2.13 | | | 0.24 + 0.43 | | |
| Temperature | 25.43 + 4.14 | | | 22.00 + 2.86 | | | 28.50 + 2.22 | | |
| Dominance rank | 3.66 + 1.72 | | | 3.55 + 1.76 | | | 3.75 + 1.67 | | |
| Age | 8.86 + 4.71 | | | 10.60 + 6.23 | | | 7.28 + 1.40 | | |
| Glide-ratio | 11.68 + 4.05 | | | 12.40 + 4.21 | | | 11.10 + 3.80 | | |
| Exit altitude from previous thermal | 463.81 + 250.78 | | | 448.13 + 241.39 | | | 477.91 + 258.33 | | |
| Distance to previous thermal | 653.11 + 932.68 | | | 683.40 + 1070.37 | | | 625.86 + 788.52 | | |

**Table S3. Outputs of models investigating the drivers thermal selection.** Estimates for continuous variables are scaled and provided with their 95% confidence intervals (95% CI), hence can be used as a dimensionless effect size. “All events” refers to the model considering decision events where available thermals previously used but potentially empty at time of choice while “Presence only” indicates the model focusing on decision events where vultures had to choose only between thermals currently used by conspecifics, with their respective sample size in brackets. Scaled variables are indicated by an asterisk.

| | *All events (N = 178)* | | | *Presence only (N = 61)* | | |
|---|---|---|---|---|---|---|
| *Predictors* | *Estimates* | *95% CI* | *p* | *Estimates* | *95% CI* | *p* |
| Distance to previous thermal * | -0.14 | [-0.40, 0.12] | 0.295 | -0.16 | [-1.42, 1.08] | 0.798 |
| Thermal already used | -0.06 | [-0.49, 0.37] | 0.784 | 0.04 | [-0.90, 0.96] | 0.931 |
| Preferred affiliate present | -0.78 | [-1.56, 0.007] | 0.052 | -0.21 | [-1.37, 1.00] | 0.732 |
| Mean social bond * | 0.02 | [-0.31, 0.35] | 0.907 | -0.46 | [-1.14, 0.24] | 0.194 |
| Maximum vertical speed * | -0.17 | [-0.40, 0.07] | 0.163 | -0.42 | [-0.87, 0.05] | 0.071 |
| Number of conspecific present * | 0.95 | [0.51, 1.38] | **<0.001** | 0.95 | [0.20, 1.68] | **0.013** |
| Hierarchy difference * | -0.001 | [-0.27, 0.27] | 0.991 | 0.12 | [-0.25, 0.48] | 0.506 |
| ** Mean + SD prior to scaling* | | | | | | |
| Distance to previous thermal | 754.81 + 1220.90 | | | 562.49 + 991.70 | | |
| Mean social bond | 0.13 + 0.09 | | | 0.13 + 0.09 | | |
| Maximum vertical speed | 2.57 + 0.94 | | | 2.42 + 0.91 | | |
| Group size | 0.73 + 0.83 | | | 1.16 + 0.79 | | |
| Hierarchy difference | - 2.34 + 27.78 | | | - 3.03 + 39.93 | | |

**Table S4. Outputs of models investigating the drivers of social information use.** Estimates for continuous variables are scaled and provided with their 95% confidence intervals (95% CI), hence can be used as a dimensionless effect size. Scaled variables are indicated by an asterisk.

| | *Threshold of 1.55 m* | | | *Threshold of 1.30 m* | | | *Threshold of 1 m* | | |
|---|---|---|---|---|---|---|---|---|---|
| *Predictors* | *Estimates* | *95% CI* | *p* | *Estimates* | *95% CI* | *p* | *Estimates* | *95% CI* | *p* |
| Distance to previous thermal * | -0.14 | [-0.40, 0.12] | 0.295 | -0.14 | [-0.40, 0.12] | 0.294 | -0.14 | [-0.40, 0.12] | 0.290 |
| Thermal already used | -0.06 | [-0.49, 0.37] | 0.784 | -0.06 | [-0.50, 0.37] | 0.770 | -0.04 | [-0.48, 0.39] | 0.833 |
| Preferred affiliate present | -0.78 | [-1.56, 0.007] | 0.052 | -0.80 | [-1.59, -0.01] | **0.046** | -0.68 | [-1.48, 0.11] | 0.093 |
| Mean social bond * | 0.02 | [-0.31, 0.35] | 0.907 | 0.03 | [-0.28, 0.35] | 0.831 | -0.04 | [-0.36, 0.28] | 0.791 |
| Maximum vertical speed * | -0.17 | [-0.40, 0.07] | 0.163 | -0.16 | [-0.40, 0.07] | 0.172 | -0.18 | [-0.42, 0.05] | 0.130 |
| Group size * | 0.95 | [0.51, 1.38] | **<0.001** | 0.95 | [0.52, 1.38] | **<0.001** | 0.92 | [0.49, 1.34] | **<0.001** |
| Hierarchy difference * | -0.001 | [-0.27, 0.27] | 0.991 | 0.003 | [-0.27, 0.27] | 0.984 | -0.003 | [-0.27, 0.27] | 0.980 |
| ** Mean + SD prior to scaling* | | | | | | | | | |
| Distance to previous thermal | 754.81 + 1220.90 | | | 754.81 + 1220.90 | | | 754.81 + 1220.90 | | |
| Mean social bond | 0.13 + 0.09 | | | 0.13 + 0.10 | | | 0.12 + 0.10 | | |
| Maximum vertical speed | 2.57 + 0.94 | | | 2.57 + 0.94 | | | 2.57 + 0.94 | | |
| Group size | 0.73 + 0.83 | | | 0.73 + 0.83 | | | 0.73 + 0.83 | | |

| | | | |
|---|---|---|---|
| Hierarchy difference | - 2.34 + 27.78 | - 2.34 + 27.78 | - 2.34 + 27.78 |
